# BCM PDX Portal: An Intuitive Web-based Tool for Patient-Derived Xenograft Collection Management, as well as Visual Integration of Clinical and Omics Data

**DOI:** 10.1101/2023.02.15.528735

**Authors:** Heidi Dowst, Apollo McOwiti, Fei Zheng, Ramakrishnan Rajaram Srinivasan, Anadulce Hernandez-Herrera, Nino Rainusso, Lisa Brubaker, Qizhi Cathy Yao, Michelle Redell, Alexandra Stevens, Seth Lerner, Sarah Woodfield, Andres F. Espinoza, John D. Landua, Susan G. Hilsenbeck, Lacey E. Dobrolecki, Michael T. Lewis

## Abstract

**Objective:** Mouse Patient-Derived Xenograft (PDX) models are essential tools for evaluating experimental therapeutics. Baylor College of Medicine (BCM) established a PDX Core to provide technical support and infrastructure for PDX-based research. To manage PDX collections effectively, de-identified patient clinical and omics data, as well as PDX-related information and omics data, must be curated and stored. Data must then be analyzed and visualized for each case. To enhance PDX collection management and data dissemination, the BCM Biomedical Informatics Core created the BCM PDX Portal (https://pdxportal.research.bcm.edu/).

**Materials and Methods:** Patient clinical data are abstracted from medical records for each PDX and stored in a central database. Annotations are reviewed by a clinician and de-identified. PDX development method and biomarker expression are annotated. DNAseq, RNAseq, and proteomics data are processed through standardized pipelines and stored. PDX gene expression (mRNA/protein), copy number alterations, and mutations can be searched in combination with clinical markers to identify models potentially useful as a PDX cohort.

**Results:** PDX collection management and PDX selection of models for drug evaluation are facilitated using the PDX Portal.

**Discussion:** To improve the translational effectiveness of PDX models, it is beneficial to use a tool that captures and displays multiple features of the patient clinical and molecular data. Selection of models for studies should be representative of the patient cohort from which they originated.

**Conclusion:** The BCM PDX Portal is a highly effective PDX collection management tool allowing data access in a visual, intuitive manner thereby enhancing the utility of PDX collections.

## BACKGROUND AND SIGNIFICANCE

Several laboratories worldwide have developed patient-derived xenograft collections representing a broad range of cancer types. Many of these individual laboratories now participate in PDX consortia, including the NCI PDXNet and EuroPDX, where data standards and infrastructure in support of PDX-based research are being developed.(1,2) While individual laboratory collections are likely to remain the norm, many of these collections are being incorporated into large repositories (e.g. the NCI Patient-Derived Models Repository (PDMR)).(3) In aggregate, these models are representative of a significant cross-section of the patient population and are generally publicly available to researchers upon request.

For PDX-based studies to be maximally useful, investigators need to identify a relevant PDX cohort in a manner similar to that which would be applied to patient selection in clinical trials. While the number and variety of models available for some organ sites is now sufficient to perform meaningful translational research, often the clinical and genomics data needed to refine the PDX study are difficult to access. Common clinical eligibility criteria categories in cancer studies may include patient demographics, age at onset of the disease, staging, patient response to prior treatments, and metastasis status, to name a few. Many of these data elements are now formalized in the Minimal Information for Patient-Derived Tumor Xenograft Models (PDX-MI), a data standard developed jointly by the NCI PDXNet and EuroPDX consortia, and adopted for use in this and other web-based tools developed by these consortia and by governmental and commercial entities (e.g. PDXFinder (https://www.pdxfinder.org/), NCI Patient-Derived Models Repository (https://pdmr.cancer.gov/), PDXNet Portal (https://portal.pdxnetwork.org/), Mouse Models of Human Cancer database (https://www.jax.org/jax-mice-and-services/in-vivo-pharmacology/oncology-services/pdx-tumors).(1-4)

## OBJECTIVE

To facilitate management of PDX collections and to address the problem of poorly accessible patient annotations for PDX study design, the Baylor College of Medicine Patient-Derived Xenograft and Advanced In Vivo Models Advanced Technology Core (BCM PDX-AIM Core)(https://www.bcm.edu/research/atc-core-labs/patient-derived-xenograft-and-advanced-in-vivo-models-core) and the Biomedical Informatics Group in the Dan L Duncan Comprehensive Cancer Center (DLDCCC), in conjunction with DLDCCC clinicians, designed and built a web portal, the BCM PDX Portal (https://pdxportal.research.bcm.edu/). Unlike other existing PDX Portals, that allow data display for PDX selection, the primary purpose of the BCM PDX Portal is to enhance PDX collection management. In addition to collection management functions, and like some other portals, the BCM PDX Portal also allows data integration, analysis, visualization, and dissemination.(5)

A primary use case of the portal is to allow researchers to select PDX models easily based on clinical and omics data, including such categories as patient biomarker status, laboratory results, treatment response, gene expression, and mutations. Cancer-specific Collection Summary pages in the portal display disease specific fields that contribute to a determination of patient treatment or inclusion in clinical trials, thus potentially increasing the clinical-translational value of the data derived using PDX models. Detailed views for each model are provided, including a visualization of a patient’s clinical timeline, to chronicle the time point of specimen donation, as well as to summarize treatments and responses that may impact PDX model behavior experimentally.

In addition to hosting BCM PDX-related data, the PDX Portal allows PDX-generating groups at any institution worldwide to host and manage their own data independently. External contributors can choose whether to make the data publicly available or to simply manage models privately. Currently, the public PDX Portal displays data representing collections at Baylor College of Medicine, Texas Children’s Hospital, and the Huntsman Cancer Institute of the University of Utah. Private collections include the University of Basel, Switzerland.

## MATERIALS AND METHODS

### Software Architecture

The BCM PDX Portal architecture follows the micro-services architecture strategy. The core services required by the PDX Portal user interface, application data, histology image viewer, genomic and copy number variation (CNV) graphs, data management services, and user authentication and authorization exist as separate software applications integrated together via an application programming interface (API). This strategy allows us to address three important concerns typical in a project involving bioinformatics and informatics components that require a heterogeneous development team to implement. First, services are developed independently and deployed by individuals possessing the relevant domain knowledge. For example, the genomics application uses genomic toolsets, while the histology image services use NoSQL based toolsets. Secondly, it allows different services to be developed using the most appropriate programming language. Lastly, services have different load requirements and usage characteristics, so there is a need for independent scalability. For example, the computation required for generation of genomics graphs is best suited for a managed cloud environment such as Amazon Web Services (AWS) or Azure, where auto-scaling occurs without any downtime.

The implementation details are as follows. The portal is composed of five separate, but interacting, applications. The PDX Portal user interface (UI) application is designed as a web portal. This is implemented as a JSF2 application, deployed on a JEE8 application server. The web UI is composed of HTML5, CSS3 and JavaScript web technologies. The portal UI components utilize Bootstrap4 UI components, ReactJs DOM framework and HighCharts web graphics API. The UI is designed to be highly responsive and thus easily viewable via various web-supporting devices. The portal web application collects information from the other portal services and aggregates these to provide the various portal UI elements. These other application services each expose a RESTful API for data exchange with the PDX Portal web UI application. The UI components are secured by OpenID, while RESTful endpoints are protected with the use of a token. The various applications are implemented with a combination of JEE8 and Spring frameworks. The genomics analysis pipeline is based on Linked Omics whose portal UI is PHP based.(6) The applications achieve persistence with a combination of a relational database solution and a NoSQL database solution.

### Data Modeling

PDX Portal relational data are stored in an Oracle 12c database schema; modeling is based on the PDX-MI data standard, which allows for the BCM PDX Portal to export data in a format that facilitates exchange with the PDXNet (https://portal.pdxnetwork.org/), the PDMR (https://pdmr.cancer.gov/), and the EuroPDX PDXFinder (http://www.pdxfinder.org/).(1-3) Major additions to the BCM PDX Portal data model include the adoption of customizable biomarkers linked to either the patient table for clinical tests or the PDX model table for laboratory tests, pediatric cancer staging systems, and additional measurements of treatment response, as seen in the entity relationship diagram provided in Appendix 1.

### Features Home Page

The focal point of the PDX Portal Home Page is a bar graph depicting the PDX collection size by disease site, a subset of which may be available for distribution to the research community (Figure 1). Dark blue columns in the graph represent publicly available PDX models, while crimson stacked columns, visible only upon login and validation of collection specific permissions, represent private models. Private models are typically those in development, obtained from elsewhere, or somehow problematic (e.g. slow growing, unusual histology, divergent gene expression versus tumor-of-origin). A collection can be selected from the graph by clicking on the interactive bar representing that collection or by use of the Collections menu bar, which redirects the user to the PDX Collection Summary page tailored specifically to a given organ site. This is the primary page available for initial cohort selection and is segmented into up to four tabs when all data is available: Patient Clinical View, Gene View, CNV View, and Mutation View tabs.

**FIGURE 1.**
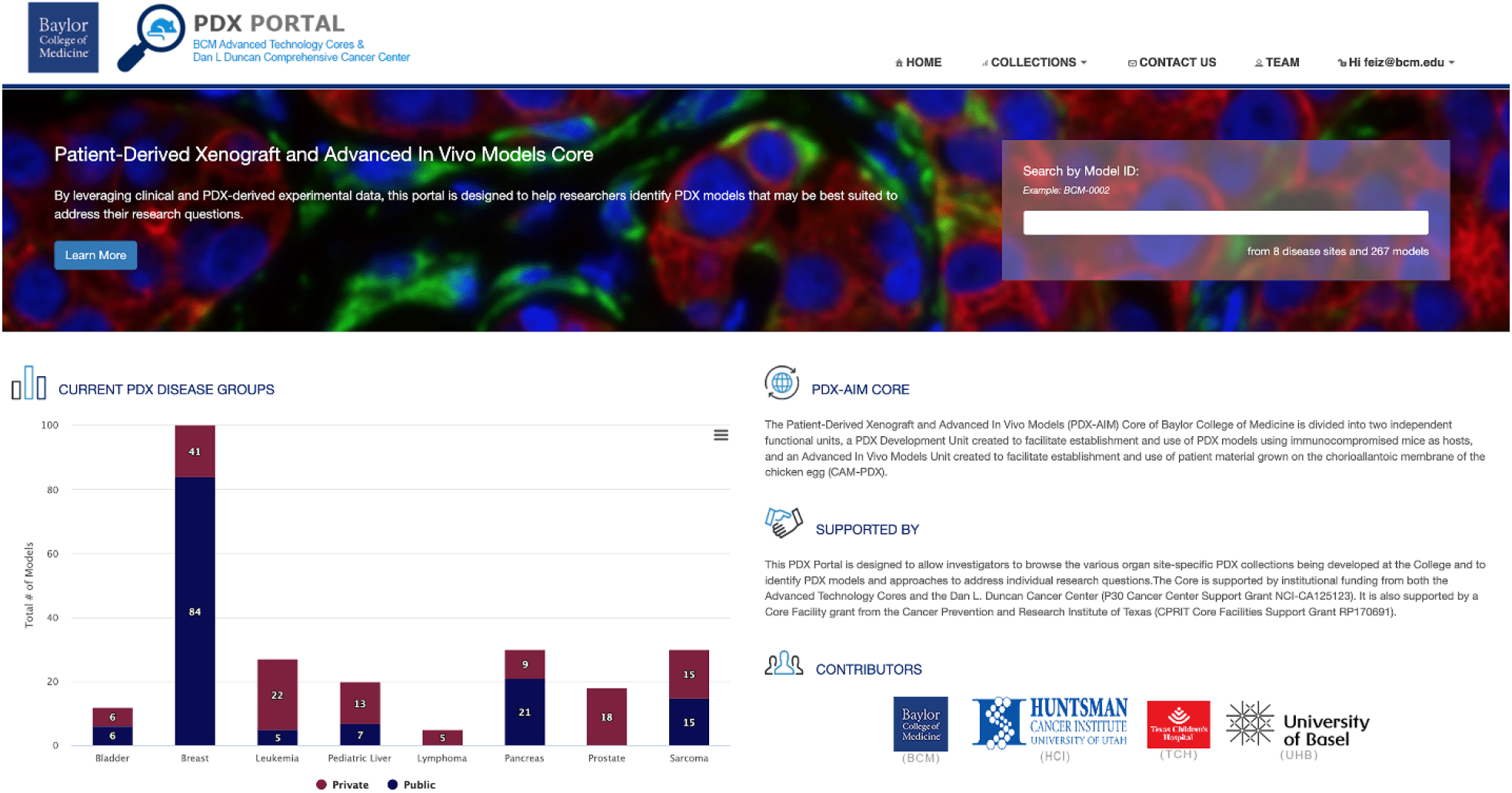
PDX Portal Homepage. Bar plot is interactive and allows for clicking on a PDX collection to view the collection details.

### Patient Clinical View

The Patient Clinical View introduces the user to the patients represented by the PDX models with a series of graphics (Figure 2). Information in this view has proven critical for appropriate study design and cohort selection for PDX-based translational research. Ideally, patient data to be shown in the PDX Portal is collected concurrently with receipt of tissue samples for transplantation.

**FIGURE 2.**
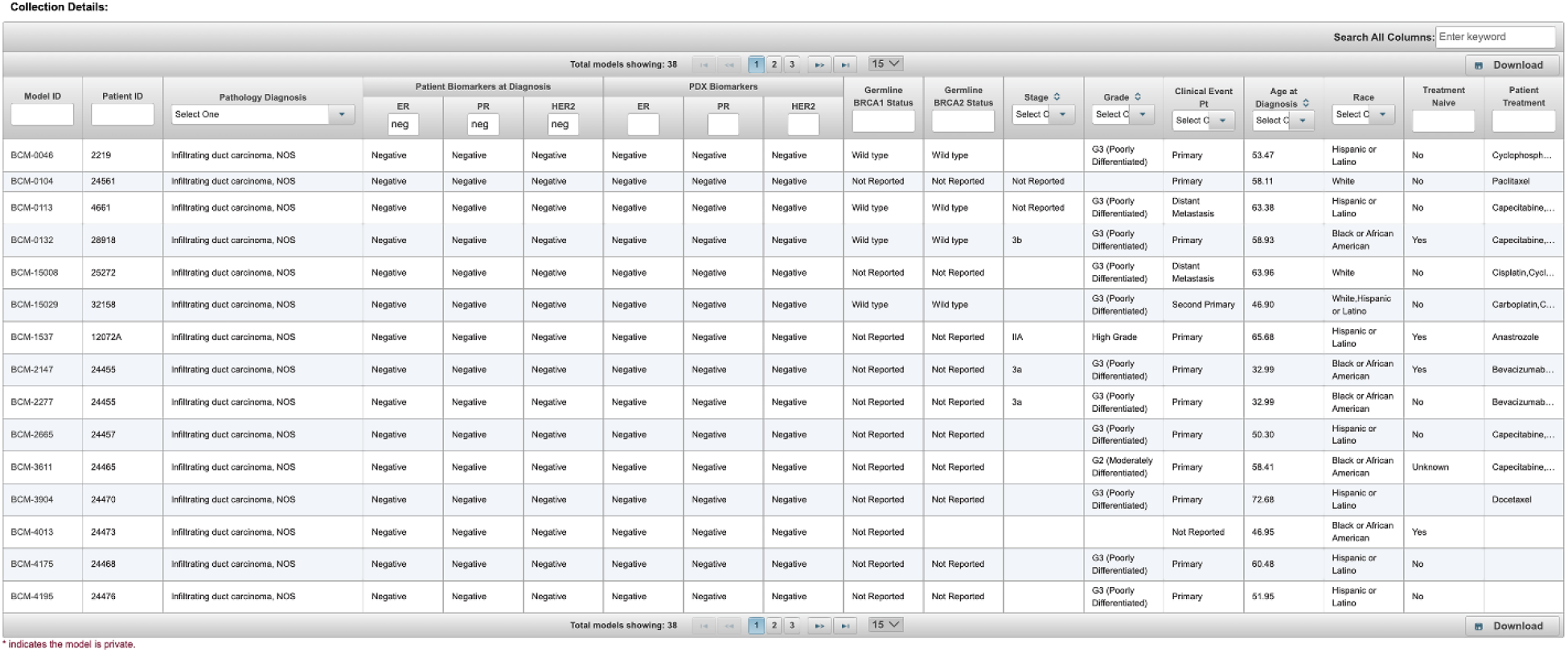
PDX and Patient Clinical Summary View. Summary graphs displaying the distribution of patients and PDX models in each disease collection. A few patient samples generated more than one model based on variations in the transplant conditions.

Many of the data elements displayed in the summary page graphics are common across all PDX organ sites such as Race and Ethnicity, Gender, Age at Diagnosis, Histology, Stage and Grade, Treatment Naive (lack of treatment prior to specimen collection), Clinical Event Point at specimen collection (Primary, Recurrence, Metastasis), and Treatments or Drugs. Other clinical data elements that are critical for the identification of a specific cohort are disease specific and range from clinically performed IHC/FISH tests such as CA 19-9 in pancreatic cancer, to blast counts performed serially on leukemia patients. The list of disease specific attributes included in the BCM PDX Portal is shown by cancer type in Table 1.

**Table 1.**
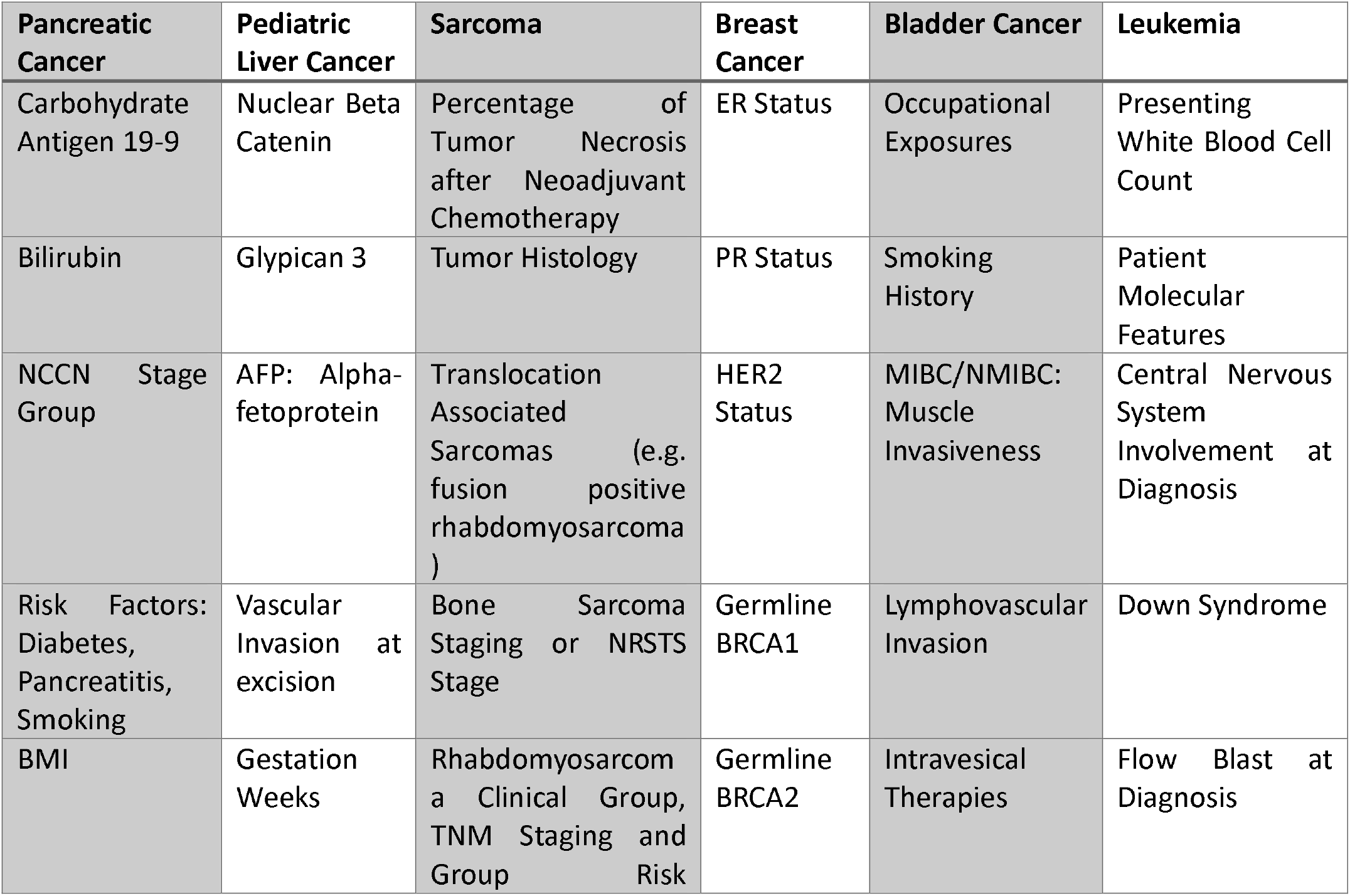
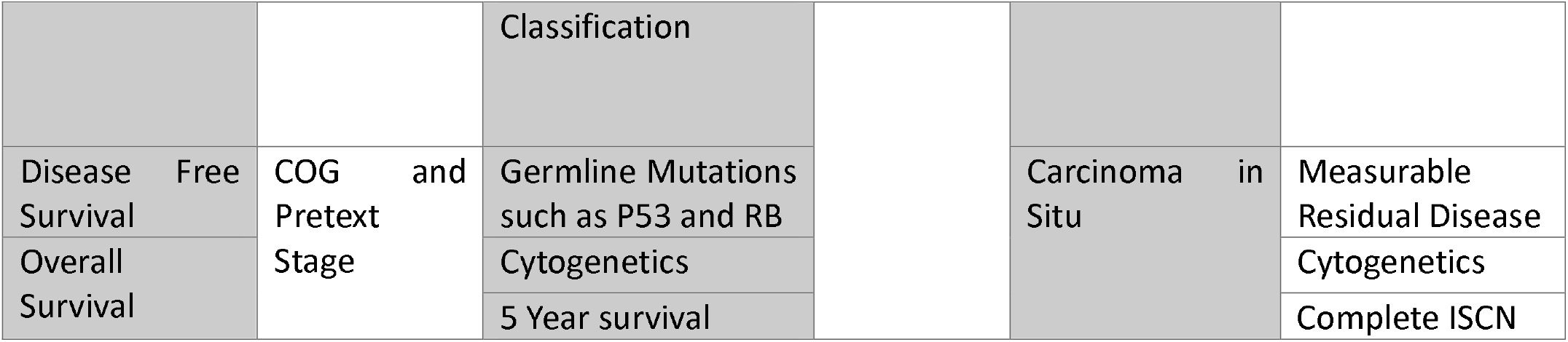
Disease Specific Clinical Attributes. Clinically relevant patient attributes which contribute to treatment decision making have been included in the PDX portal to enable model selection based real-world evidence.

Other summary graphs include patient risk factors or co-morbidities such as pancreatitis, diabetes, and smoking history (Figure 2).(7) Attributes for model selection include clinically-tested patient germline or somatic mutations that have an established disease linkage such as germline BRCA1 or BRCA2 (breast and ovarian cancer), as well as status for other clinically-relevant biomarkers (e.g. immunostaining for the estrogen and progesterone steroid hormone receptors and for ErbB2 (HER2) amplification and/or overexpression in the breast cancer PDX collection). Additional summary data may indicate environmental circumstances, such as premature birth as described on the pediatric liver PDX collection page reflecting the effect of NICU treatments to developing immature livers.(8) These disease specific data elements are real world tools used by oncologists when making treatment recommendations in the clinical setting.

Treatment response is another attribute included, whenever available, that incorporates defined data elements for each PDX collection. While many solid cancers use pathologist assessments, magnetic resonance imaging (MRI), computed tomography (CT), or positron emission tomography (PET) scans to measure change in tumor size in response to treatment and to estimate Residual Cancer Burden (RCB), other cancers such as leukemia and lymphoma employ other markers as indicators of response as shown in Table 2. For the Leukemia collection, the PDX Portal reports the Measurable Residual Disease (MRD) value as an indicator of patient treatment response while for the Osteosarcoma collection, the PDX Portal reports the percentage of tumor necrosis from surgical excision of the primary tumor after neoadjuvant chemotherapy.(9-11) In the Texas Children’s Hospital pediatric liver collection, Alpha-fetoprotein (AFP), a serial clinical measurement, is an early indicator of patient treatment response and is also displayed in the PDX Portal as model search criteria on the respective collection summary page.(12)

**Table 2.** Disease-Specific Measurements of Treatment Response. Clinical evaluation of patient response to treatment is measured by a variety of evidence and clinical standards based on the cancer diagnosis. The PDX portal has incorporated the variety of treatment measurements found in table 2.

| Disease Specific PDX Collection | Treatment Response Measurements |
| --- | --- |
| Solid tumor cancers | RECIST Response |
|  | Pathologic Response |
| Pancreatic cancer | Changes in CA 19-9 Levels |
| Leukemia | Measurable Residual Disease/Flow Blast |
|  | Cerebral Spinal Fluid Involvement |
| Osteosarcoma | Good Response (>90% of tumor necrosis) |
|  | Poor Response (<90% of tumor necrosis) |
| Pediatric liver cancer | Changes in Alpha-fetoprotein (AFP) Levels |

The Collection Details table found at the bottom of the collection summary page, which displays a data matrix researchers can use to view available models in the selected collection, can be used to filter models by each of the general and specific disease markers. Searching and filtering can be applied to individual columns in this table at the top of each column while a global search field and download button are also provided on the upper right-hand corner of the table as seen in Figure 3. Models meeting multiple clinical criteria are selected by filtering in each desired column.

**FIGURE 3.**
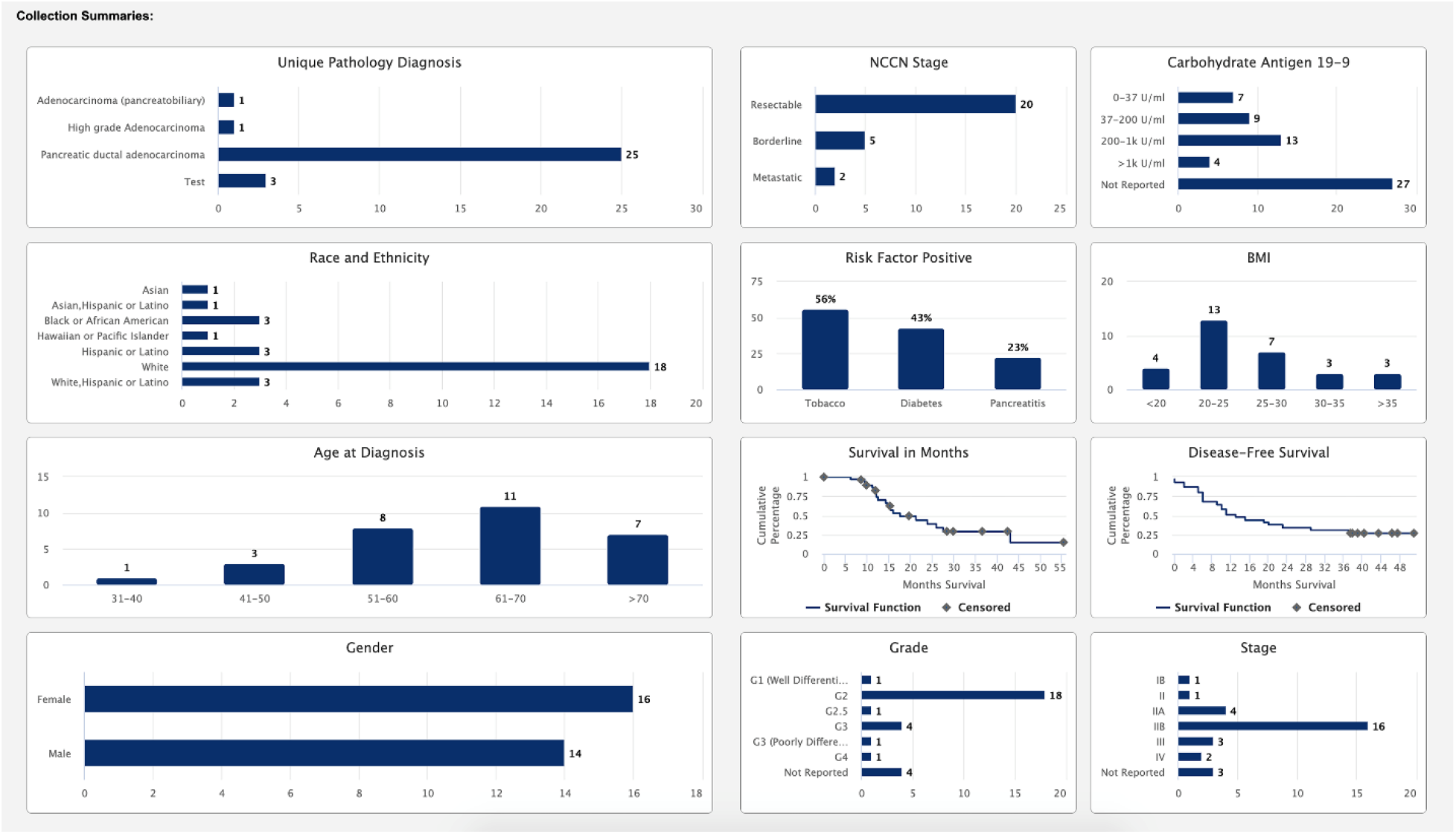
Collection Details Table. Summary collection data table which allows for filtering by multiple data variables to identify models of interest.

### Gene View, CNV View, and Mutation View

The Gene View, CNV View and Mutation View tabs allow for visualizing various omics data types across samples. Minimally, mRNA expression by RNAseq or microarray should be included with each collection. However, the portal is flexible and can display any quantitative omics data type including proteomics and metabolomics. Each view has options to search for genes by HGNC symbols, aliases, and gene name descriptions or to upload lists of HGNC symbols. The Gene Expression view additionally enables creating a sub-plot to drill down on samples that match user-specified clinical biomarker criteria, visualizing trends that differ between the overall dataset and a clinical subset of interest as shown in Figure 4.

**FIGURE 4.**
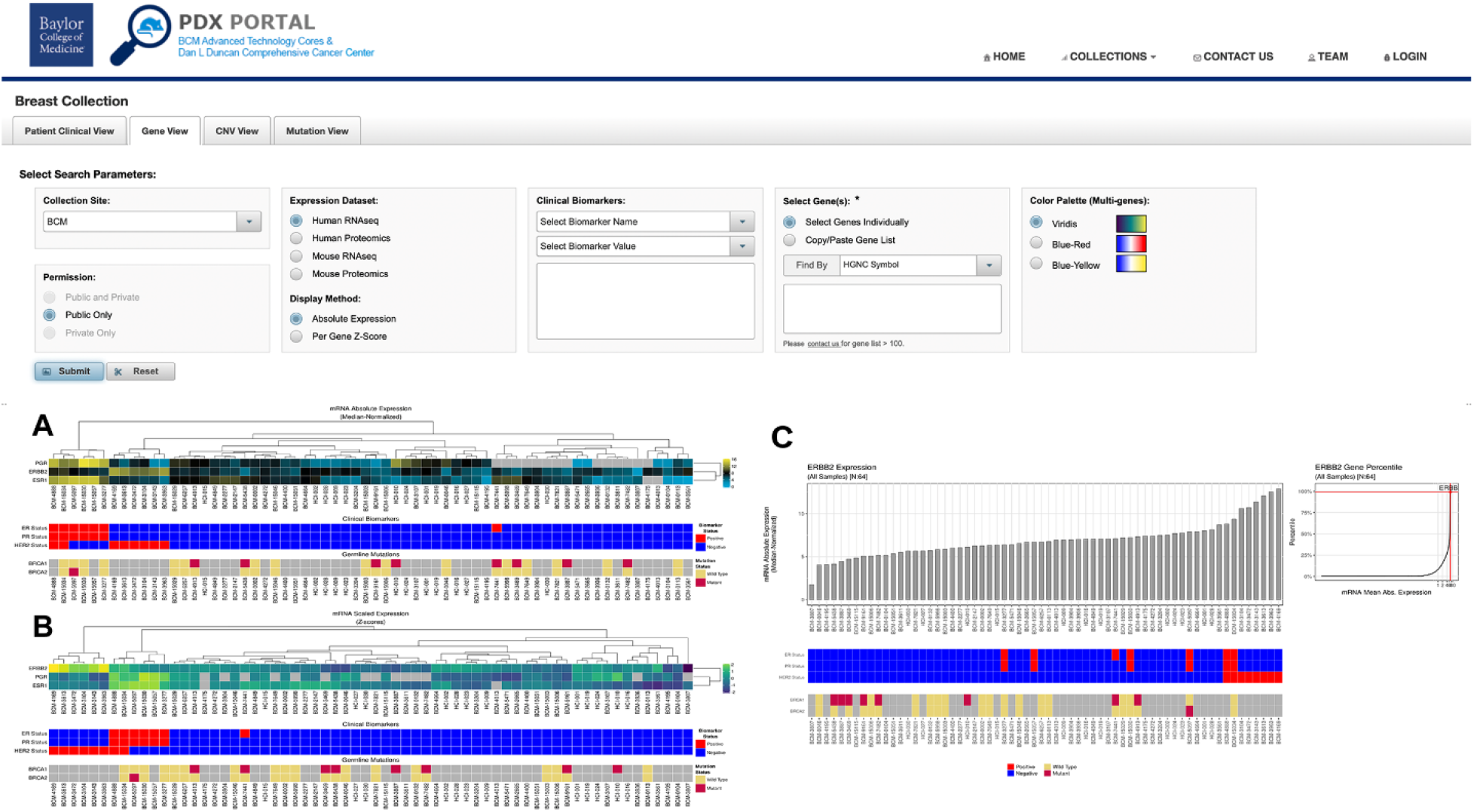
Gene Expression View: Top Panel) Search and display parameters; A) Multi-gene heatmap (absolute expression), B) Multi-gene heatmap (per-gene Z-scores); C.) Single gene expression bar graph

Because mouse and human mRNA or protein can be distinguished from one another computationally, the Gene Expression view (Gene View) provides options to visualize expression levels in either the PDX tumor or the mouse host.(13,14) Expression of two or more genes can be displayed in a heat map using multiple color palettes: viridis for colorblind individuals, red-blue, or yellow-blue (Figure 4, top panel). For two or more genes, data can be plotted using either “absolute expression”, which allows comparison of gene expression levels both within a sample and across PDX models (Figure 4A), or a “per gene Z-score” that displays relativistic expression for a given gene across samples (Figure 4B). Gene expression level for single genes is displayed in a bar graph (Figure 4C). Both methods have utility for model selection and data interpretation.

The CNV View tab affords methods to visualize calculated copy number alterations at the gene level. These alterations can be visualized in terms of the relative intensity of the alteration compared to its surrounding genomic region (Figure 5A) or the type of alteration, such as deletion, loss, gain, and amplification (Figure 5B). These CNV plots can then be correlated with transcript, or protein expression levels.

**FIGURE 5.**
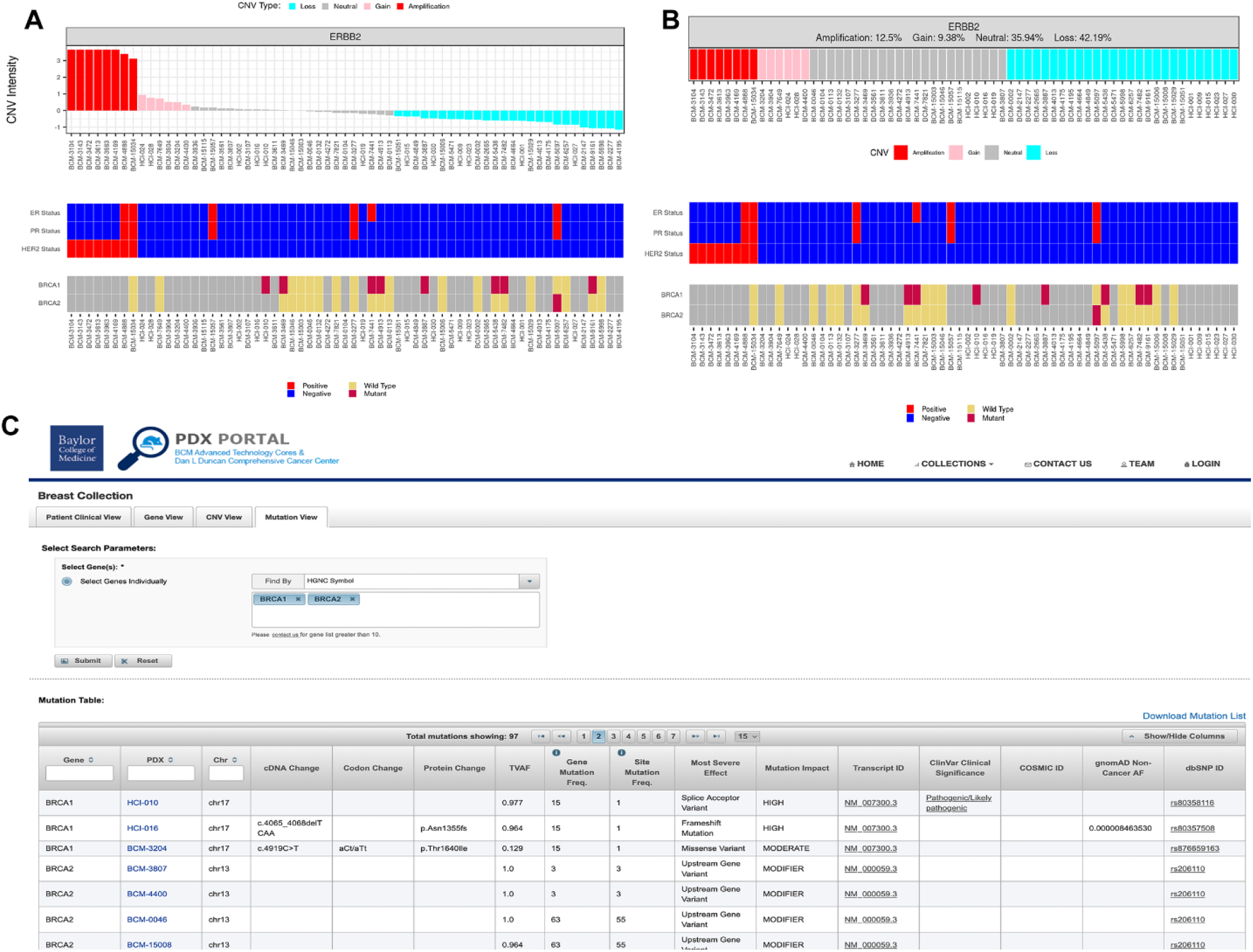
Copy Number and Mutation Views: A) CNV View (intensity plot); B) CNV View (type plot); C.) Mutation View

The Mutation View generates a mutation table for one or more genes of interest that includes the PDX model identification (hyperlinked to the Model Detail Page), chromosomal location, cDNA change relative to a reference sequence, codon change, protein change (amino acid substitution), the variant allele frequency (VAF), the frequency of the alteration in the PDX collection, and mutation impact (Figure 5C). Hyperlinks to outside databases such as dbSNP, COSMIC, and Clinvar are also provided in the results table to annotate the gene mutations.

### Model Detail Page

Moving back to the Collection Details table at the bottom of the Patient Clinical View page, an individual model can be selected by double clicking on the model name, after which a specific Model Details page loads. The Model Details page is also segmented by tabs: Patient, PDX Model, Histology, Metastasis, and Patient Treatment. The Model Details page provides additional details specific to the individual model selected from the collection summary page.

The focus of the Patient tab is the visualization of the clinical timeline of the patient beginning at cancer diagnosis. The timeline contains procedures, treatments, responses, and follow-up, when available (Figure 6). The patient clinical procedures are displayed in purple and are labeled with chronological event ID numbers. Rolling the cursor over these procedure events displays information about the event, along with patient age at event, disease progression, staging, and diagnosis, if applicable. Rolling the cursor over the treatment events, colored in teal, displays the treatment regimen or drug received along with the age at start, age at stop of treatment, duration of treatment, and the tumor’s response to the treatment, if available.

**FIGURE 6.**
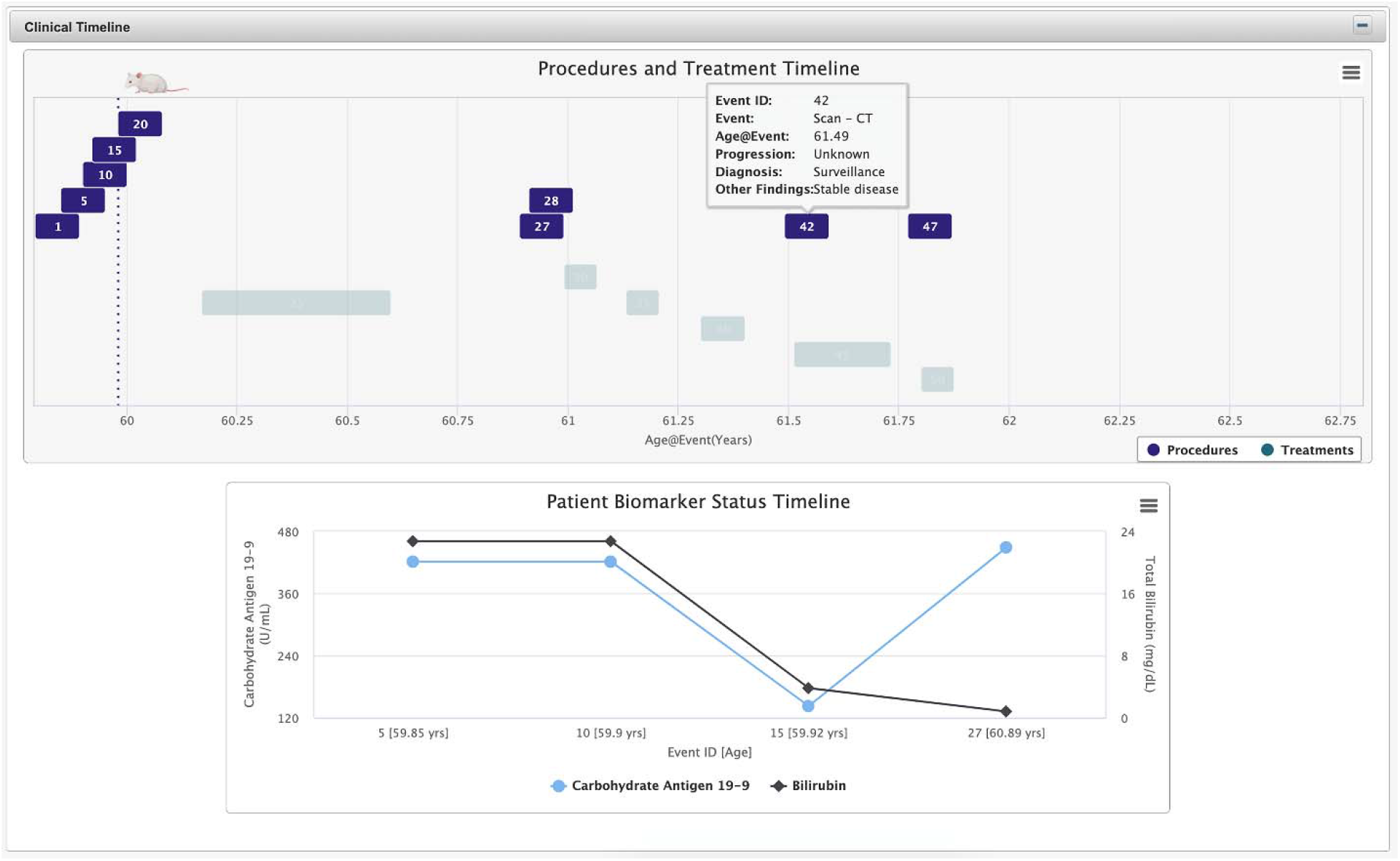
Patient Clinical Timeline. Clinical events relating to the patient’s history of cancer are show on the timeline along with treatment events which detail treatment response when available.

A critical element of the timeline is the mouse icon, which identifies the time point at which the specimen for PDX generation was collected. Where multiple models were developed from the same patient, each model is designated by a separate mouse icon. Rolling the cursor over the mouse icon on the Clinical Timeline graph displays the official name of the PDX model. The clinical timeline quickly identifies drugs that the patient received prior to specimen collection, and indicators of the PDX model treatment response to drugs that the patient subsequently received after collection. Below the clinical timeline, the line graph, or bar graph, depending on the collection, displays serial patient marker expression over time. In a pancreatic cancer example (Figure 6) the graphic indicates a drop in the CA 19-9 marker at event 10, followed by an increase at event 15, possibly indicative of a response, followed by a resurgence of the cancer in the patient.(15)

The PDX Model tab displays transplantation conditions for PDX development, as well as individual model mutations and copy number variations, information useful for experimental planning if specific tumor types are required (e.g. ESR1-positive). Another important feature of the BCM PDX Portal is the ability for a researcher to evaluate the correlations between the PDX model and the patient. One feature that enables this is the Metastasis tab, which displays known patient metastatic sites side-by-side with organ sites evaluated in the PDX model. Similarly, the Histology tab displays images of hematoxylin/eosin-stained tissue sections along with a selection of immunohistochemistry (IHC) images from the patient and PDX model specimen side-by-side. From these images, correlations between the histology of the PDX tumor and the patient’s tumor-of-origin can be evaluated.

The final tab, Patient Treatment, is a chronological list of drugs received by the patient along with the clinical response, pathologic response, and reason for stopping treatment, whenever available. Unfortunately, these data are some of the more difficult clinical data to obtain. Often preferred indicators of patient response are unavailable in the electronic health record (EHR), making it critical for the data model to accept multiple indicators. The “Reason Stopped” field, although not collected in a structured format in medical records, can provide insights into treatment side effects that may have resulted in an incomplete course of treatment or worsening symptoms that that may indicate disease progression. Since many of these patients were not enrolled in clinical trials, these may encompass non-categorized adverse events. This additional treatment column allows for recording information outside of the standard treatment criteria represented by RECIST 1.1 or College of American Pathology (CAP) pathologic response values.

## RESULTS

The BCM PDX Portal is designed to facilitate translational research using PDX models and is available for public use. The portal differentiates itself from other PDX tools by providing PDX collection management tools, as well as by providing integrated omics analysis and display functionality. When models of interest are identified on the BCM PDX Portal, contact information for the program lead for each organ site can be found on the Contact Us page. Models can then be requested via a Material Transfer Agreement from the institution of record. For BCM models, investigators may use. Once the MTA is executed, PDX models can be shipped either fresh (if available) or viably frozen for re-transplantation.

The following PDX study use cases illustrate the use of the PDX Portal in study design. One such completed study on triple negative (ER-, PR-, HER2-) breast cancer used nine PDX models in which amplification of chromosome 12p was found to associate with emergence of docetaxel resistance and with carboplatin sensitivity.(16) Although the work for this study was conducted prior to construction of the BCM PDX Portal and without the benefit of fully abstracted clinical records, the use case illustrates how model selection could have been accomplished more efficiently using the BCM PDX portal. This study would have commenced by selecting the Breast Cancer PDX collection from the PDX Portal home page and filtering the Collection Details table for “negative” in the ER, PR, and HER2 columns. Additionally, filtering on docetaxel in the Patient Treatment column identifies 24 models where the patient received docetaxel as part of their treatment regimen. Further review of the Model Details page and the Patient Treatment tab for the recorded patient responses to docetaxel would have been used as a possible predictive measure. Viewing the patient clinical timeline also provides insights into continuous drug exposure or companion drugs such as carboplatin where responsiveness was also investigated in this article. As a follow-up to the study, the PDX Portal could be used further to identify those lines with BRCA1/2 mutations (or any other) as well as specific copy number variations for these models.

Another powerful use of PDX models is to aid in the evaluation of novel therapeutic agents for efficacy in cancer treatment. Model selection for these types of studies can be done in a variety of ways. Models can be selected randomly, which has proven less productive, or models can be selected rationally based on the expression level of the drug target or other predictive biomarker (Figure 4) or on the presence or absence of a genetic mutation (Figure 5C). Using the PDX Portal, investigators can quickly identify models with increased or decreased gene expression, or harbor a mutation of interest. Conversely, model selection can be based off of a specific clinical attribute of the patient.

To highlight the utility of such an approach, a search could be performed to identify models that might be sensitive to an anti-TEM8 (Anthrax Receptor 1, ANTXR1) CAR-T cell treatment regimen. This would be accomplished by mining the PDX Portal RNAseq data to find models expressing high levels of ANTRX1 at the RNA and/or protein level. In this particular study, models expressing high levels of ANTRX1 RNA were evaluated for their level of tumor vascularity since anti-TEM8 therapy targets tumor-associated vascular endothelium and blocks neovascularization. Two models were selected for treatment and both showed a reduction in tumor volume when treated with the CAR-T cell therapy, with one showing a decrease in vascularization.(17)

Drug validation studies often require identification of tumors likely to and not likely to respond (negative controls) based on the expression level of the target of interest. In a study of an ERN1 (IRE1) inhibitor (ERN1 is a downstream target of MYC), RNAseq data were leveraged to determine the expression level of MYC in TNBC PDX models. On the Gene View tab of the portal, the Clinical Biomarkers were set to “negative” for ER, PR, and HER2, then the MYC gene was selected and the search submitted. Models with varying levels of MYC mRNA were selected, and MYC protein expression levels were validated subsequently by IHC staining. Four models with lower expression of MYC (predicted non-responders or poor responders) and two models with higher expression of MYC (predicted responders) were then treated with the inhibitor. As the level of MYC expression increased across models, the response to the inhibitor increased, consistent with their likelihood of MYC dependency.(18)

## DISCUSSION

These published examples highlight the utility of the BCM PDX Portal to facilitate PDX-related translational research. As more PDX models and their associated clinical- and PDX-related data are incorporated into the site, opportunities will present themselves to enhance ongoing research activities and open new research questions. For instance, data from the Gene View tab in the breast collection are currently being mined to find additional PDX models that express genes in the Hedgehog pathway for evaluation of their role in EMT and breast cancer metastasis.(19) Functionality to identify pre- and post-treatment PDX pairs or primary and metastatic PDX pairs will also aid in the elucidation of chemo-resistance and metastasis mechanisms, respectively.(20) Finally, given the racially and ethnically diverse patient populations that Baylor College of Medicine, Texas Children’s Hospital, and other institutions served, models in these collections will be useful for study of tumor-intrinsic racial disparities, as models derived from Hispanic or Latino and African American patients can be easily identified and compared with those derived from Caucasian patients.

## CONCLUSION

The BCM PDX Portal currently displays information for six of the twelve active cancer type-specific PDX programs at BCM/TCH: bladder, breast, and pancreas, as well as pediatric liver, leukemia, and sarcoma. Eventually, all twelve PDX collections will be incorporated. These additional cancer types include ovarian, prostate, glioblastoma, lung, head and neck, as well as pediatric brain cancers. The portal will continue to incorporate disease specific markers as they are identified, as well as candidate response indicators given that differential expression of selected genes has already been demonstrated to be a valuable resource for model selection in PDX-based preclinical studies. That said, the BCM PDX portal is already capable of hosting data from anywhere in the world for PDX collection management and data display, with data controlled virtually entirely by the contributors themselves.

The BCM PDX Portal represents one of the leading examples of adoption of existing PDX data modeling standards, where they exist, and establishing new data recommendations based on real world clinical events. Time and effort put into promoting the adoption of data standards for PDX models will facilitate inter-institutional PDX modeling and data sharing further. As collaborations and model sharing occur, model annotations will continue to grow as additional studies increase the knowledgebase for each model and PDX models in general. Finally, the BCM PDX Portal has been integrated seamlessly with other web-based resources under development (e.g. the Molecular and Imaging Response Analysis of Co-Clinical Trials (MIRACCL) resource (https://miraccl.research.bcm.edu/)).

## ACKNOWLEDGEMENTS

This work was supported, in part, by the Dan L Duncan Cancer Center P30 Cancer Center Support Grant CA125123, an NCI PDXNet PDX Trial Center Cooperative Agreement U54 CA224076, a Co-clinical Imaging Research Resource Program (CIRP) grant U24 CA226110, as well as a Core Facility Support grants (RP170691 and RP220646) from the Cancer Prevention and Research Institute of Texas (CPRIT).

The authors wish to thank Ping Gong, Christina M. Sallas, and Alaina N. Lewis for PDX model data entry.

The contents do not represent the views of the U.S. Department of Veterans Affairs or the United States Government.

## AUTHOR CONTRIBUTIONS

Conceptualization: MTL, HD

Software: AM, FZ, RRS, HD

Validation: LD, JL, AHH

Resources: LD, JL, AHH

Formal Analysis: SGH, RRS

Writing – OD: HD

Writing – Review & Editing: MTL, LD

Data Curation: AHH, LB, NR, QCY, MR, AS, SW, SL, AE

Visualization: LB, NR, QCY, MR, AS, SW, SL, AE

Supervision: MTL, HD, SGH

Funding Acquisition: MTL

## APPENDIX I

**APPENDIX 1.**
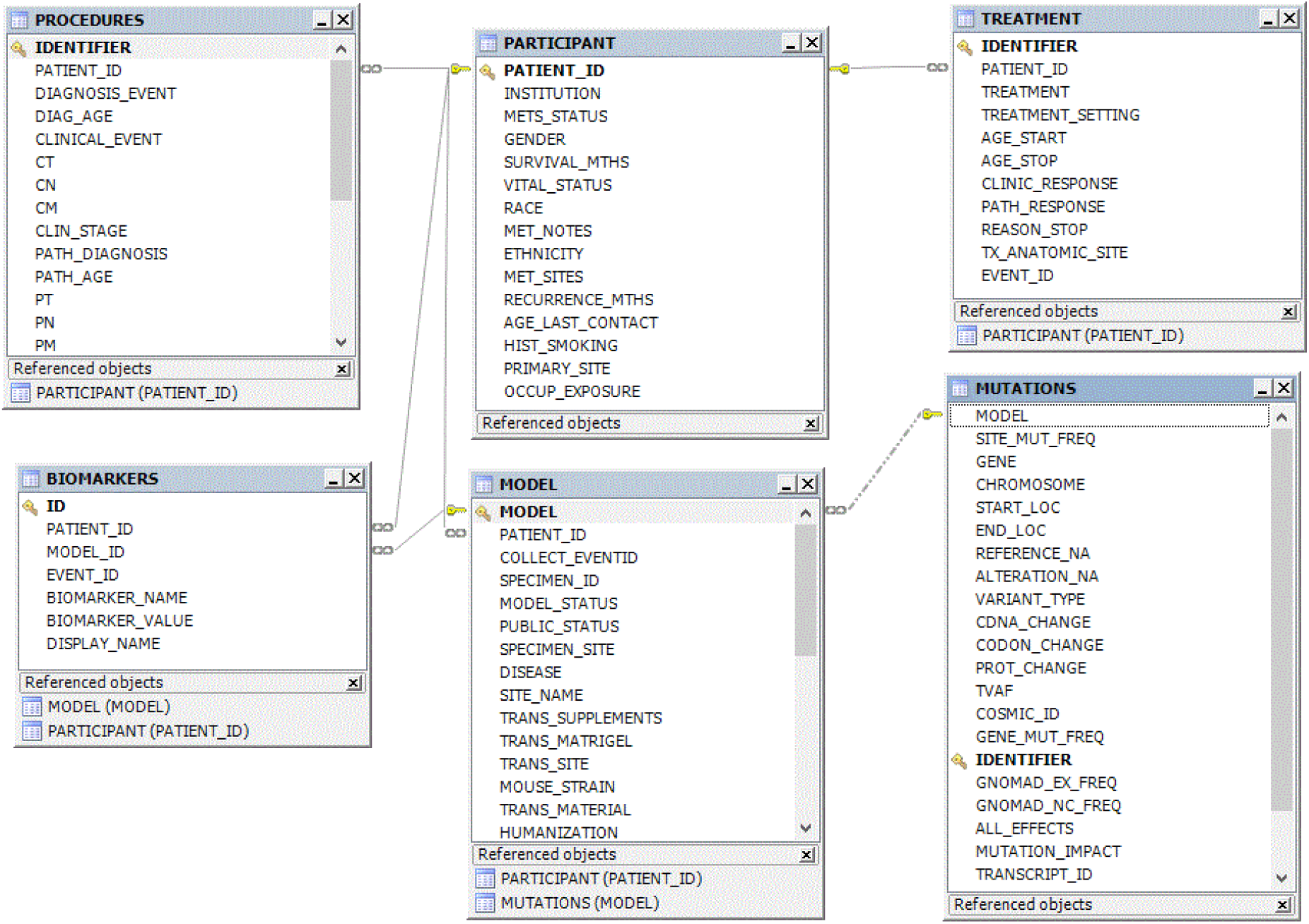
PDX Portal Entity Relationship Diagram: While the complete data model contains additional tables which enable application operations, the essential data is represented by the six tables shown in this Entity Relationship Diagram. The Patient table, containing patient demographics and risk factors, is the parent table with a foreign key to the Model table. These two tables are both linked by foreign keys to the biomarker table, listing all unique patient and model biomarkers. The patient table is linked by a foreign key in a one-to-many relationship to the Procedures and Treatments tables which enables chronological tracking of the patient clinical timeline. The Mutations table containing all gene annotations is linked by a foreign key to model table.

